# 2-D organoids demonstrate specificity in the interactions of parasitic nematodes and their secreted products at the basal or apical intestinal epithelium

**DOI:** 10.1101/2024.12.16.628471

**Authors:** Ruby White, Jose Roberto Bermúdez-Barrientos, Elaine Robertson, María A. Duque Correa, Amy H. Buck

## Abstract

The intestinal epithelium is a selectively permeable barrier that absorbs water and nutrients, senses the luminal environment, and orchestrates submucosal immune responses. Some gastrointestinal nematodes occupy both submucosal and luminal niches and can cause profound changes in epithelial cells. Yet, there is little understanding of the specificity of parasite-host interactions with epithelial cells and the differential responses at the apical or basal sides. Organoids enable uncoupling of parasite-driven responses from immune-driven effects on the epithelium, and two-dimensional (2-D) organoids allow for controlled access to both sides. Here we used 2-D enteroids to model *in vivo* localisation of *Heligmosomoides bakeri* (*H. bakeri*) fourth-larval (L4) and adult stages and identify localisation-specific effects on gene expression. Using bulk RNA sequencing, we found an enhanced upregulation of IFN-stimulated genes (ISGs) during basal co-culture and an overall stronger response to L4 parasites than adults. Further, *H. bakeri* live worms induced much stronger effects on gene expression than their isolated excretory-secretory (ES), but cells that internalised *H. bakeri* extracellular vesicles (EVs) displayed up-regulation of ISGs, repair genes, and YAP/TAZ signalling. Our results demonstrate that nematodes and their products, including EVs, modify the host epithelium in a localisation-dependent and cell-specific manner.

## INTRODUCTION

The intestinal epithelium is a critical mucosal barrier responsible for water and nutrient uptake and protection of the body from the external environment while maintaining homeostasis ^1,2^. This barrier consists of a semi-permeable mucous layer, and an underlying polarised epithelial layer containing multiple cell types, including absorptive enterocytes and secretory enteroendocrine, tuft, Paneth, and goblet cells, which are self-renewed by intestinal stem cells (ISCs) ^1,2^. The polarisation of the intestinal epithelium underpins its many critical functions with the apical and basal epithelium having different biological roles: the former is specialised in selective absorption of water and nutrients, and the latter is involved in integrating signals to and from the rest of the body ^3,4^. In particular, the intestinal epithelium is integral to the clearance of gastrointestinal (GI) nematode infections by sensing parasites, initiating immune responses and executing effector functions such as increased mucus production and peristalsis referred to as the ‘weep and sweep’ response ^5^. These effector functions are a result of drastic remodelling of the host intestinal epithelium as a response to parasite-induced tissue damage, parasite excreted/secreted (ES) products, or host immune factors produced to control infection e.g. Interleukin(IL)-4 and IL-13 ^6^.

*Heligmosomoides bakeri* (*H. bakeri*) is a rodent infective GI nematode that resides in either the submucosa or lumen of the small intestine during the different stages of its life cycle ^7,8^. Granulomas of host immune cells including myeloid cells, CD4^+^ T cells, and NK cells form around encysted fourth-stage (L4) larvae approximately four days post-infection (p.i.) ^9–15^. In parallel, ISCs in the epithelium overlying granulomas display a loss of the canonical ISC marker *Lgr5* coupled with an appearance of cells transiently expressing Sca1 (*Ly6a*) and other genes associated with foetal epithelium, which are suggested to represent a repair response to tissue damage during larval invasion ^13,16^. ISC expression of foetal genes, but not the loss of *Lgr5* expression, was shown to be dependent on Interferon (IFN)-γ signalling, produced by CD8^+^ T and NK cells ^13,15^. Expansion and activation of tuft and goblet cells also occurs during early infection, with tuft cells responding to *H. bakeri* by secreting IL-25, which leads to the activation of Innate Lymphoid Cells (ILC) type 2 (ILC2s) that in turn produce IL-13 resulting in further tuft and goblet cell expansion ^17–21^. Despite extensive knowledge on the effects that the immune response to parasites has on the epithelium, we still have little understanding of the direct modifications induced by *H. bakeri* parasites or their ES products on the intestinal epithelium.

GI nematodes produce a heterogenous milieu of ES products including small molecules such as metabolites and ascarosides as well as proteins and nucleic acids that are released individually or within extracellular vesicles (EVs) ^20–24^. We previously reported that *H. bakeri* EVs are internalised by the small intestinal MODE-K epithelial cell line, which displays features resembling enterocytes. The EVs reduced expression of *Dusp1* and the IL-33 receptor subunit encoding gene *Il1rl1* within MODE-K cells ^22^. However, it is not clear how these observations extrapolate to the intestinal epithelium as epithelial cell lines do not replicate the physiology and diversity of epithelial cell populations *in vivo.* 3-D organoid cultures more closely replicate the cellular composition and structural features of the intestinal epithelium, allowing investigation of direct interactions of live parasites and their ES products with the epithelium in absence of the immune system ^6,25^. For instance, while there is an overall increase in tuft cell numbers during *H. bakeri* infection, adult *H. bakeri* ES (HES) treatment of 3-D organoids surprisingly abrogated IL-4/IL-13 induced tuft cell hyperplasia ^26^. Additionally, whilst the aforementioned foetal stem cell signature induced by *H. bakeri* infection was suggested to be dependent on IFN-γ, HES treatment of 3-D organoid cultures has been shown to elicit this transcriptional profile on a distinct cluster of cells, termed revival stem cells (revSCs), independently of immune cell factors ^16^. These examples demonstrate that organoids enable the differentiation of changes to the host epithelium driven by the parasite from those triggered by the host immune response. However, these studies have focused on the impact of adult HES delivered to the basal epithelium of 3-D organoids. Given that immune sensing receptors are known to localise to the basal epithelium, we reason it may be more responsive to parasites and their ES products, not reflecting host-parasite interactions that take place at the apical side *in vivo* ^27^.

Here, we use a 2-D small intestine organoid (enteroid) model that recapitulates key features of the host epithelium to test these hypotheses and investigate the host epithelial response to *H. bakeri* L4 larvae on basal epithelium, versus adult worms and their ES products at the apical surface. We found basal co-culture with either L4 larvae or adult parasites elicits a strong transcriptional response with upregulation of interferon-stimulated genes (ISGs) and repair genes, many of which are targets of the YAP/TAZ transcription factors involved in tissue repair ^28^. Apical stimulation with adult *H. bakeri* showed fewer transcriptional changes than basal treatment suggesting interactions at the apical membrane result in more subtle host responses. We then queried the relative contributions of EVs and non-vesicular HES at the apical epithelium and found weaker effects compared to live parasites. However, we show that only a specific subpopulation of cells internalised *H. bakeri* EVs and comparison of EV+ and EV-cells revealed a striking upregulation of genes associated with the IFN and repair responses in cells that internalised the vesicles. These findings demonstrate the direct and localisation-specific modulation of the epithelium by *H. bakeri* in the absence of host immune factors and highlight a previously unappreciated link between *H. bakeri* EVs and epithelial repair during infection.

## Results

### 2-D enteroids mimic apical and basal *in vitro* niches of *H. bakeri* and respond to parasites in a life-stage and localisation-dependent manner

2-D organoid methods are well established for mouse caecal and colonic epithelia ^29,30^. However, current methods for murine 2-D enteroid cultures either do not establish confluent monolayers ^31,32^ or do not report on the presence of dividing, absorptive and secretory cellular populations of the intestinal epithelium ^33,34^. Here, we modified existing protocols (Fig 1a) to establish confluent monolayer cultures of mouse enteroids containing the diversity of cellular populations found in the small intestine (Fig 1b - i). Our 2-D enteroid cultures demonstrated some degree of spatial organisation as previously reported for caecaloids ^30^ (Supplemental Movie 1). Specifically, they contained Ki67^+^ proliferative cells (stem and transit-amplifying (TA) cells), which self-organised into proliferative loci (Fig 1c & e), surrounded by non-proliferative zones containing enterocytes, Paneth, tuft, goblet and enteroendocrine cells (Figures 1c-i). Cellular polarisation was observed in our 2-D cultures with the brush border at the apical surface identified by the colocalisation of Villin and F-actin (Fig 1d).

**Figure 1:**
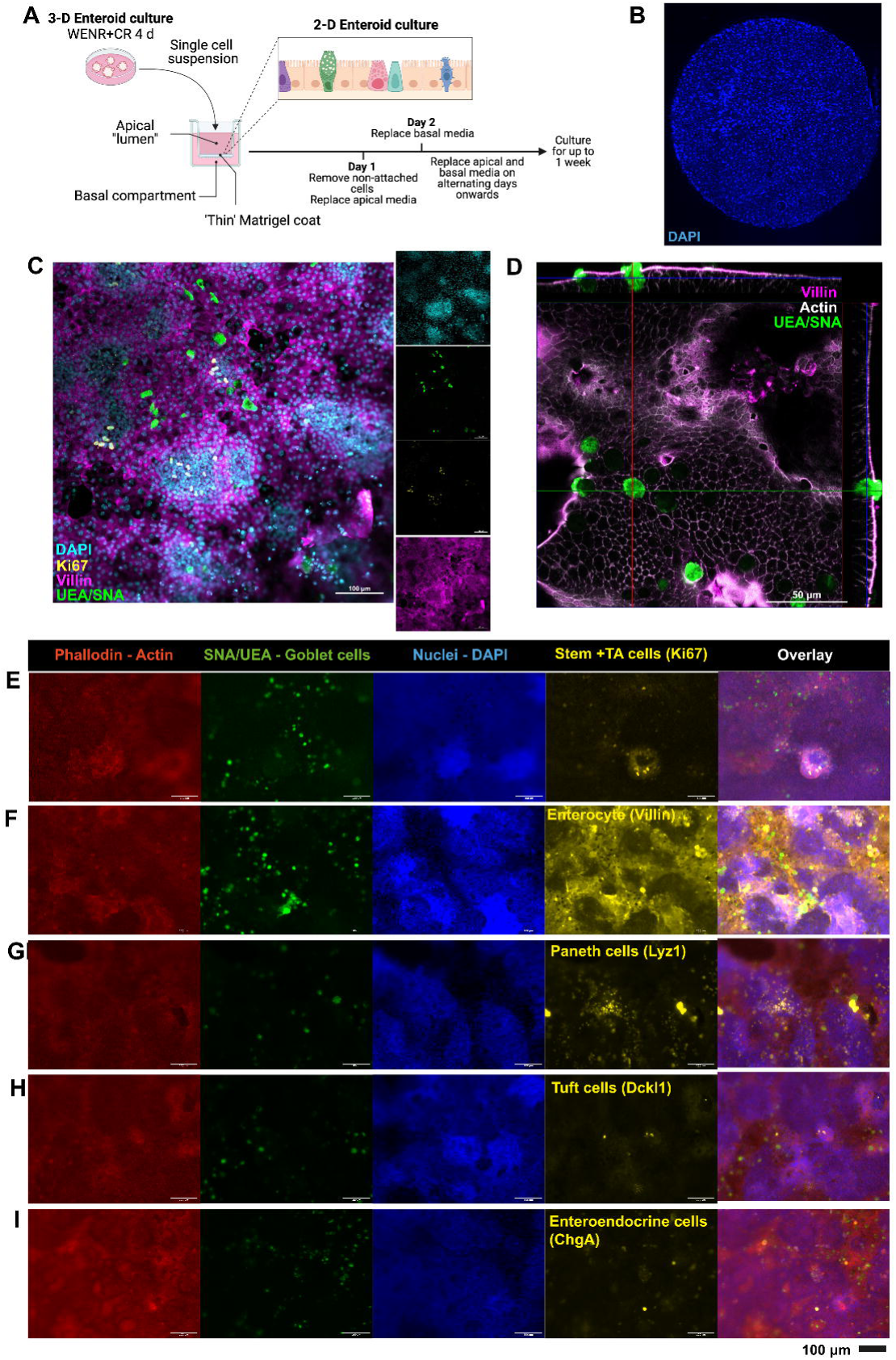
2-D enteroids recapitulate the cellular composition and spatial organisation of the small intestinal epithelium. A) Schematic of experimental culture conditions for the generation of 2-D enteroid cultures in Matrigel-coated transwell plates. B) Representative fluorescence microscopy image showing complete transwell coverage by 2-D enteroid monolayer stained with 4’,6-diamindino-2-phenylindole (DAPI) for nuclei (blue). C) Representative confocal immunofluorescence image showing organisation of Ki67+ Alexa Fluor 647 (AF647, yellow) cells into proliferative loci, surrounded by terminally differentiated goblet cells labelled with UEA & SNA FITC (green). Nuclei are stained with DAPI (cyan) and F-actin labelled with phalloidin (Magenta). D) Z-stack projection of confocal immunofluorescence image demonstrating apical brush border of epithelial cells by co-staining of microvilli marker Villin AF647 (Magenta) and F-actin using Phallodin AF594 (White). *Ulex europaeus agglutinin* I (UEA) fluorescein (FITC) and *Sambucus nigra* Lectin (SNA) FITC binds lectins present in mucus of goblet cells, UEA & SNA FITC (green) can be seen emerging through the brush border. E-I) Representative fluorescence microscopy images of 2-D enteroids with DAPI-stained nuclei, phalloidin AF594 binding F-actin, and UEA/SNA labelling goblet cells, and antibody-based immunohistochemistry in AF647 of E) villin on microvilli of enterocytes, F) the Ki67+ stem and transit-amplifying cells, G) lysozyme 1-expressing Paneth cells, H) Dckl1 expression on tuft cells and I) chromogranin A on enteroendocrine cells.

Next, we used our 2-D enteroids to determine the direct impact of *H. bakeri* L4 larvae or adult parasites at their respective sites in infection, either in contact with the basal or apical membranes of the epithelium. We further tested whether the life-stage of the parasite impacts the host response at the basal epithelium by comparing culture with L4 larvae or adult parasites (Fig 2a). RNA sequencing (RNAseq) analysis showed that L4 larvae co-cultured at the basal epithelium induced stronger transcriptional changes (432 up-regulated and 808 down-regulated differentially expressed genes (DEGs)) compared to co-culture of adult parasites (337 up-regulated, 718 down-regulated DEGs) (Fig 2b, Supplemental Table 1). In contrast, we observed fewer DEGs when adult *H. bakeri* was cultured with the apical epithelium (142 up-regulated, 252 down-regulated) compared to the responses of either parasite stage cultured at the basal epithelium (Fig 2b, Supplemental Table 1). Notably, basal co-cultures specifically induced stronger upregulation of several ISGs, compared to apical co-cultures (Fig 2c). Other pathways were upregulated in all co-cultures to varying degrees. For example, upregulation of epithelial repair genes such as annexins and lymphocyte antigen-6 family genes (Ly6) was evident in L4 larvae basal co-culture and adult apical co-culture, and to a lesser extent in adult basal co-culture consistent with previous studies (Fig 2c)^13,16^. Many of the genes upregulated across all co-cultures are known targets of YAP/TAZ activation (transcription factors of the hippo signalling pathway involved in intestinal epithelial repair ^28,35,36^), which were most strongly upregulated in response to L4 larvae compared to basal or apical culture with adult parasites (Fig 2c, Supplemental Fig 1). Genes downregulated across all comparisons included those related to stemness and cell cycle ^13,16,26^. To understand whether parasite co-culture impacts the differentiation of the epithelium into secretory or absorptive lineages, we performed a pathway enrichment analysis method (CAMERA) of gene profiles associated with stem, TA, secretory progenitors, Paneth, goblet, tuft, enteroendocrine and enterocyte cells from published single-cell RNAseq datasets and determined enrichment or depletion of cell types ^16,37,38^. We found that stem cells were downregulated by basal co-culture with adult or L4 larvae, and secretory progenitor cells by co-cultures of L4 larvae with the basal epithelium and adults at the apical side (Fig 2d). Of the secretory lineages, Paneth cells were downregulated by all co-culture conditions, and enteroendocrine cells showed a downregulation only after basal parasite co-cultures (Fig 2d). In contrast, enterocytes were consistently upregulated suggesting that *H. bakeri* co-culture skews differentiation of the epithelium towards the absorptive lineage (Fig 2d). Finally, revSCs were strongly upregulated in all co-culture conditions (Fig 2d). Taken together, we observed a direct response of the intestinal epithelium to *H. bakeri* parasites, which is characterised by upregulation of ISGs and repair genes and downregulation of stemness. In parallel, *H. bakeri* co-culture skewed differentiation of the epithelium towards the absorptive lineage and away from secretory cells. The magnitude of these responses was dictated by the life-stage of the parasites and its localisation within the epithelium.

**Figure 2:**
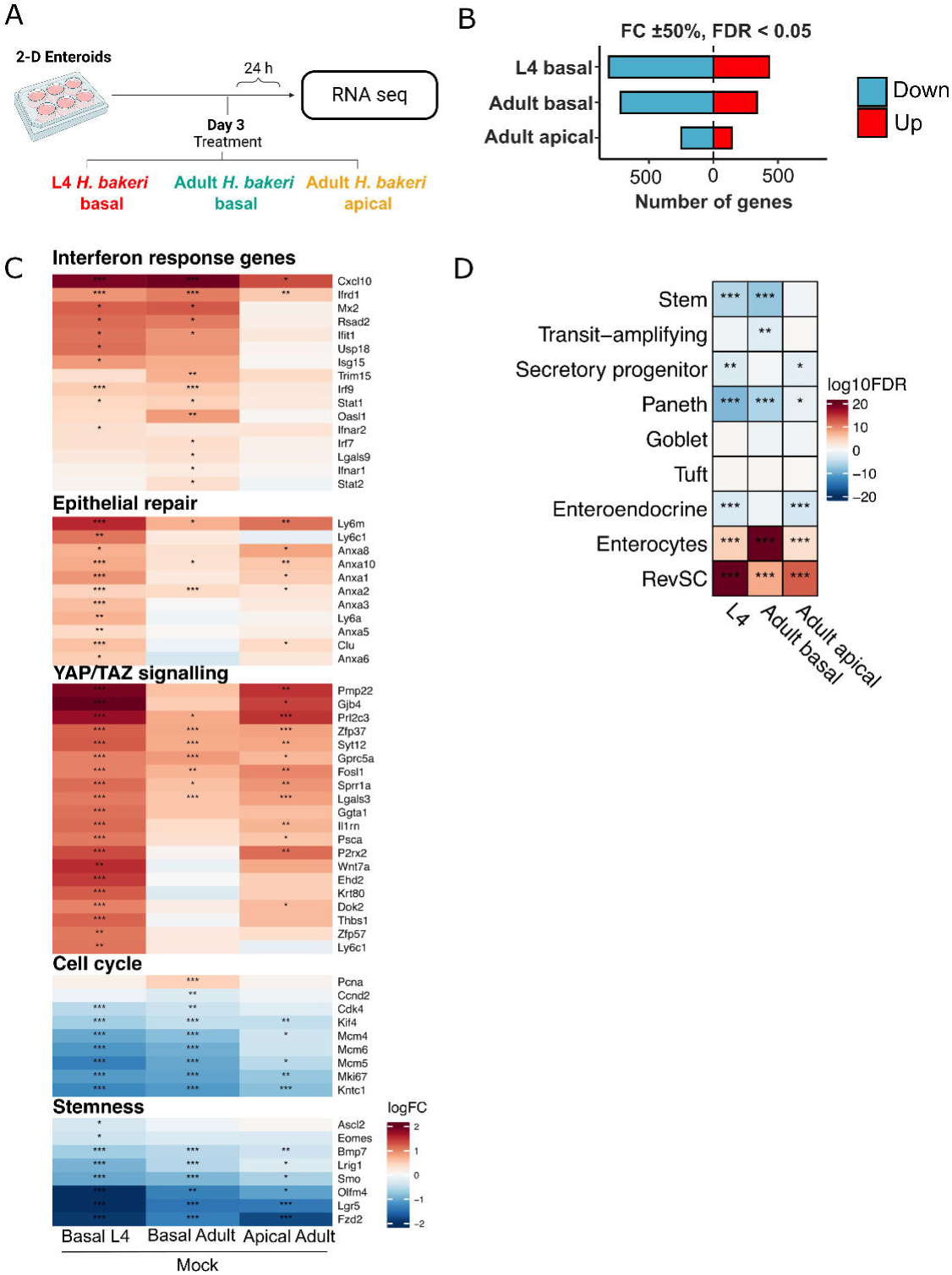
The intestinal epithelium responds in a life-stage and localisation-dependent manner to *H. bakeri* parasites. A) Experimental workflow to evaluate gene expression changes using RNAseq of 2-D enteroids treated on day 3 of culture with 10 live L4 larvae on the basal compartment, 10 adult parasites either on the apical or basal compartment, or Mock control for 24 h. B) Number of differentially expressed genes (DEGs) with a false discovery rate (FDR) < 0.05, and log2FC > 0.585 in response to L4 *H. bakeri* apical, adult *H. bakeri* apical or basal treated 2-D enteroids compared to mock. C) Heatmaps of selected differentially expressed genes (DEGs) (log2FC) functionally related to the interferon response, epithelial repair & the foetal epithelium, YAP/TAZ signalling, cell cycle and stemness in *H. bakeri* apical vs. mock, Adult basal *H. bakeri* vs. mock or Adult apical *H. bakeri* vs. mock comparisons. D) CAMERA analysis of gene sets of intestinal epithelial cell types from differential expression analysis (DEA) from 2-D enteroids co-cultured with basal *H. bakeri* L4 larvae, or basal *H. bakeri* adults, or apical *H. bakeri* adults. Cell type specific gene sets from published scRNAseq ^16,37^ . A-D) FDR * < 0.05, ** < 0.01, *** < 0.001.

### Apical co-culture with *H. bakeri* adult parasites induces stronger host responses than their ES products

Previous studies have investigated the direct responses of the intestinal epithelium to HES using 3-D organoids via basal delivery. 2-D organoids enable the delivery of live adult parasites and their ES products at the apical epithelium, mimicking *in vivo* localisation. Thus, taking advantage of this culture system, we examined the differences in the apical epithelium responses to co-culture with *H. bakeri* adults, HES depleted of EVs (EVdepHES), or EVs. Strikingly, we found that while co-culture with ten *H. bakeri* adults resulted in considerable gene expression changes in the epithelia (142 up-regulated, 248 down-regulated DEGs), the exposure to EVdepHES only induced 16 and downregulated 11 genes (Fig 3a-b, Supplemental Table 1). Further, we found no significant transcriptomic changes in the 2-D enteroid cultures treated with *H. bakeri* EVs for 24 h (Fig 3a-b). EVdepHES caused upregulation of several YAP/TAZ target genes (*P2rx2*, *Fst*, *F3*), cellular adhesion genes (*Fn1, Cldn4*) and the ISG *Trim15*, which were also upregulated by co-culture with adults. While we found similar host genes changing in response to the co-culture of live parasites and treatment with isolated EVdepHES at the apical epithelium, the former resulted in markedly stronger responses even though the dose of EVdepHES is much higher than what is expected to be derived from ten worms (Methods). Genes downregulated after EVdepHES were also consistent with those downregulated by adult co-culture, including *Mcm4* and *Kntc1* related to the cell cycle (Fig 3c). The remaining DEGs upon EVdepHES treatment did not group into a particular functional category. For example, while we observed upregulation of *Clca1,* a gene involved in mucus production, no other DEGs related to this function were detected (Supplemental Table 1).

**Figure 3:**
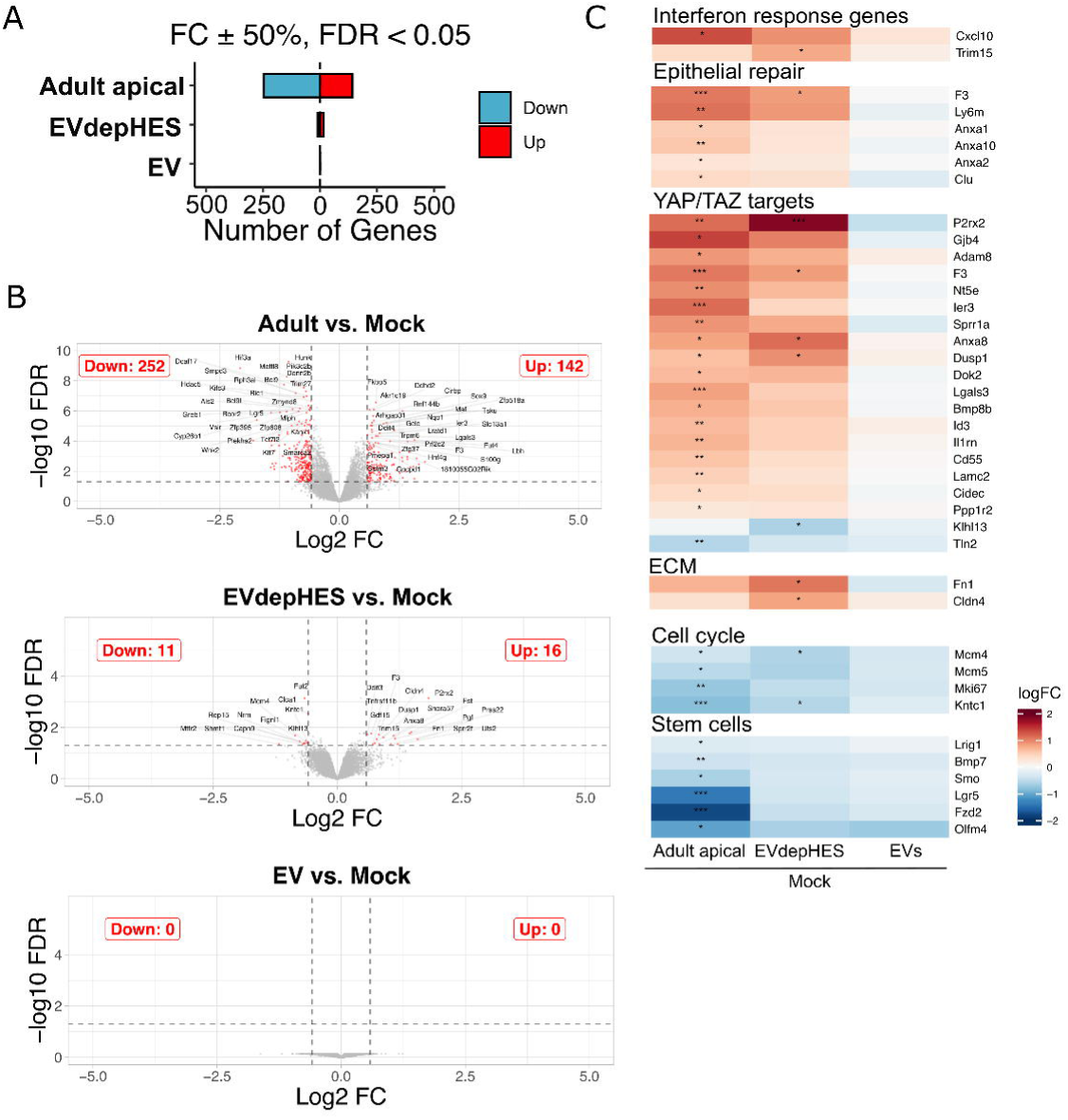
Apical co-culture of adult *H. bakeri* induces stronger host response than isolated ES products. 2-D enteroid apical co-culture with adult *H. bakeri* (10 worms), extracellular vesicles (EVs) (20 μg/ml; or 3.6×10^10^ particles/ml), or EV depleted HES (EVdepHES) (20 μg/ml), or Mock control A) Differentially expressed genes (DEGs) with FDR < 0.05, log2FC > 0.585 in each comparison, up-regulated (red), down-regulated (blue). B) mRNA expression profiles (volcano plots) from RNA-seq of adult apical co-culture (top), EVdepHES (middle), EVs (bottom) vs. Mock. Dashed lines represent FDR <0.05 and Log2FC of > 0.585 respectively. C) Heatmaps of selected differentially expressed genes (DEGs) functionally related to interferon response, epithelial repair, YAP/TAZ targets, extracellular matrix (ECM), uncategorised genes differentially expressed in EVdepHES vs. mock, cell cycle and stemness in adult apical, or EVdepHES, or EVs vs. mock.

### Enteroid cells that uptake *H. bakeri* EVs display strong induction of ISGs and wound repair genes

Given the low response of organoids to EVs at the RNA level, we aimed to understand what percentage of cells internalise EVs. We fluorescently labelled EVs with Alexa Fluor 647 (EV-647) and treated 2-D enteroids with EVs by co-culture at the apical surface for 24 h. The extent of internalisation was then quantified by flow cytometry. Control culture media was subject to an identical isolation and labelling process as EVs to account for any dye carryover (referred to as “Mock”). Importantly, 2-D enteroid cultures were dissociated using trypsin which would cleave any external EVs bound at the cell surface; therefore, the signal detected should only reflect internalised EVs. We found that 2-D enteroids treated with EV-647 displayed a significant shift in AF647 fluorescence, compared to either naïve or mock, indicating EV internalisation (Fig 4a). Strikingly, only a small percentage (x̄ = 6.4%) of total cells internalised EVs (Fig 4b). EV internalisation into cells was further validated by super-resolution microscopy (Supplemental Fig 2).

**Figure 4:**
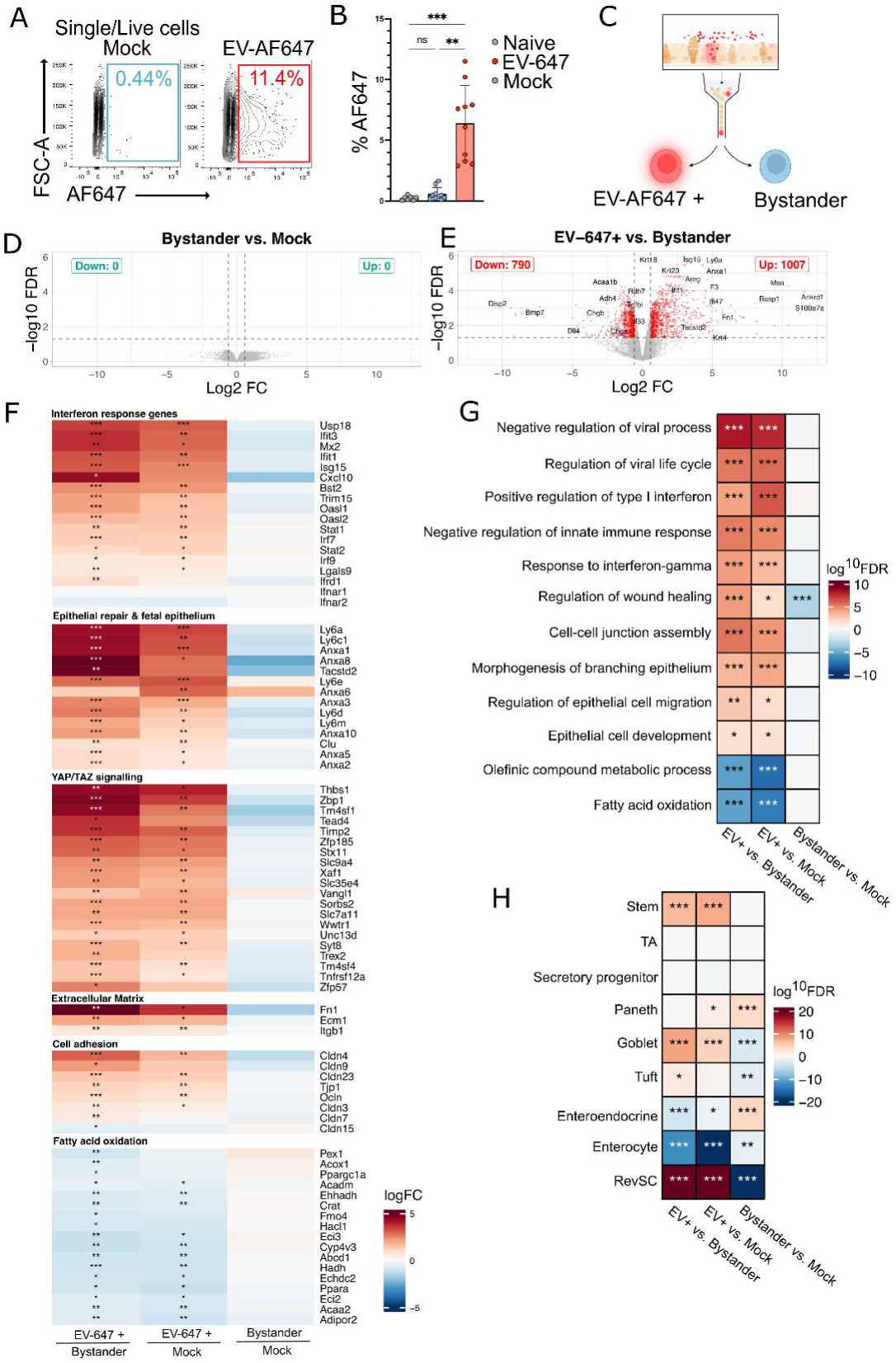
Enteroid cells which internalise *H. bakeri* EV-647 display specific induction of ISGs and wound repair genes. 2-D enteroids were co-cultured with fluorescently labelled EVs with Alexa Fluor 647 (EV-647), or a mock control for 24 h before dissociating for fluorescence assisted cell sorting (FACS) A) Representative gating of AF647 positive live single cells in Mock-647 (left) or EV-647 treated (right) treated 2D-enteroids. B) Proportion of single live cells AF647 positive indicating *H. bakeri* EV-647 uptake in untreated (grey), or mock-647 (blue), or EV-647 20 μg/ml (red) treated 2-D enteroids. C) Schematic of FACS sorted EV-647+ cells and EV-647-cells (Bystander) from 2-D enteroids co-cultured at the apical surface with EV-647. D-E) mRNA expression profile (volcano plots) of D) EV-647 vs bystander and E) Bystander vs. mock genes in red defined as DEGs. Dashed lines at FDR <0.05 or Log2FC > 0.585 F) Heatmaps of selected differentially expressed genes (DEGs) functionally related to interferon response, epithelial repair, YAP/TAZ signalling, extracellular matrix (ECM), cell adhesion and fatty acid oxidation in EV-647+ vs. Bystander, EV-647+ vs. Mock, or Bystander vs. Mock G-H) CAMERA analysis of differential expression analysis from EV-647+ enteroid cells vs. Bystander, or Mock treated cells; or Bystander vs. mock treated cells in G) selected enriched GO terms H) gene sets of cell types of the intestinal epithelium from previously published single cell RNA sequencing ^16,37^. A-B) represents 3 independent experiments, *n* = 7-10, error bars represent median and interquartile range, statistical analysis using Kruskal-Wallis with Dunn’s multiple comparisons test. \*\**P* < 0.01, \*\*\**P* < 0.001, ns > 0.05. F-H) Represents 3 technical replicates from 1 enteroid line. FDR * < 0.05, ** < 0.01, *** < 0.001.

Next, we determined the gene expression profile of cells that internalised *H. bakeri* EVs by performing RNAseq on sorted EV-647+ cells and AF647-cells from the same 2-D enteroid, or sorted cells from mock treated cells (Mock) (Fig 4c). Mock and bystander cells displayed very similar transcriptional profiles with no significant DEGs, suggesting no paracrine impact of EV treatment on bystander cells at 24 h (Fig 4d). In contrast, sorted EV-647+ cells demonstrated a striking uptake-specific transcriptional profile with 790 downregulated and 1007 upregulated genes compared to bystander cells (Fig 4 e&f, Supplemental Table 2). The transcriptome of EV-647+ cells was characterised by the upregulation of several ISGs (i.e. *Isg15*, *Mx2*, *Bst2*) (Fig 4f). We also observed an upregulation of genes related to epithelial repair, including genes that are highly expressed in the foetal epithelium and expressed in revSCs (*Ly6a*, *Tacstd2*, *Clu*, Annexins), accompanied by YAP/TAZ target genes; genes involved in extracellular matrix remodelling (*Fn1*, *Ecm*, *Itgb1*); and genes encoding tight junction proteins important for cell adhesion and barrier function (Claudins, *Ocln*); (Fig 4f, Supplemental Fig 3a). Pathway enrichment analysis supported the upregulation of both an IFN response/viral response and pathways related to wound repair and remodelling (Fig 4g). Downregulated pathways were predominantly related to metabolic alterations and specifically fatty acid oxidation, whose impairment has been recently linked to blocked differentiation of the airway epithelium (Fig 4g) ^39^. We again used CAMERA analysis to identify enrichment or depletion of cell type specific genes in the EV-AF647+ population to understand if EVs are either uptaken by specific cell types and/or play a role in differentiation towards certain cell types. Sorted EV-647+ cells were enriched in genes expressed by stem, goblet-, tuft- and most notably revSC cells and depleted in genes identified in enteroendocrine cells and enterocytes (Fig 4h). Notably, the opposite was found when comparing Bystander cells to mock treated cells: where genes expressed by RevSC, and tuft-, goblet-cells are downregulated and those expressed by enteroendocrine cells were upregulated. These data are consistent with a model where sorting of EV-647+ cells selectively removes or increases specific cell types within the bystander pool, or alternatively EV uptake is associated with differentiation towards certain lineages (Fig. 4h). In summary, while *H. bakeri* EVs are internalised by only a small proportion of epithelial cells of 2-D enteroids, these cells display a specific transcriptional program of ISGs and epithelial repair genes and gene sets expressed in stem cells as well as goblet and tuft cells. These findings suggest specificity in EV uptake and scope for functional effects in driving or regulating epithelial repair during infection.

## Discussion

Here, using 2-D enteroids we demonstrate that the life-stage and localisation of nematode parasites to the apical or basal membranes of the epithelium impacts the host epithelial response. Apical co-culture of adult parasites or their products induced a more subtle response in the epithelium compared to basal co-culture of adults, while basal L4 larvae compared to basal adults induced even larger changes. These results are in line with biological differences in the apical and basal epithelium that affect responsiveness to enteric pathogens. For example, some receptors involved in sensing PAMPs either localise to the apical or basal surface, or trigger different signalling pathways depending on the surface that encounters the ligands ^27,40,41^ .

We also observed that co-culture of 2-D enteroids with L4 larvae resulted in a stronger host response than co-culture with adult parasites, suggesting that there are life-stage-specific effects potentially resulting from the secretion of different ES products ^42^. We found that L4 larvae especially drive the upregulation of epithelial repair genes recapitulating findings *in vivo.* During *H. bakeri* infection the ISC niche is modified to a state reminiscent of the primitive foetal epithelium that is suggested to represent a repair phenotype. RevSCs arise during larval stage infection (d6p.i.) and share overlapping features with crypt cells responding to non-infectious models of epithelial damage such as irradiation, targeted ablation of Lgr5+ stem cells, and DSS colitis ^28,35,43^. These features include the expression of foetal-associated genes (*Ly6a*, *Anxa1*, *Tacstd2I*), the suppression of canonical Lgr5^+^ stem cell-associated genes (*Lgr5*, *Lrig1*, *Olfm4*, *Axin2*) and the induction of YAP/TAZ signalling and their target genes (*Ankd1*, *Ereg)* ^13,28,35,43^. Modification of the ISC niche *in vivo* has been shown to be dependent on IFN-γ, which is produced after exposure to the microbiota (this is speculated to be caused by epithelial barrier injury during migration of parasites) ^13,15^. We identified a strong upregulation of these genes in response to co-cultures with either *H. bakeri* L4 larvae, adult parasites or their ES products. Our data suggest that outwidth the presence of microbiota and IFN-γ signalling, these responses are initiated directly by the parasite or by their ES products and likely enhanced during infection by the host immune response ^13^. Surprisingly, in basal conditions where parasites were physically separated, thus removing any physical contact to the epithelium, these effects were the strongest suggesting physical contact is not essential for this host response, but may exacerbate it *in vivo*.

2-D enteroids also revealed a striking difference in the response of the apical epithelium to ten adult parasites compared to their concentrated ES products (that we estimate would be derived from >20-fold more parasites) ^42^. These results may be explained as a consequence of additional stimuli introduced by parasite co-culture that are not captured by ES treatment, for example, physical interaction, competition for the availability of nutrients in the culture media, and the release of small molecules typically excluded from ES during the isolation process such as metabolites. In this regard, we note that our purification of EVs and EV-depleted HES is more stringent than in other studies, since it used both ultracentrifugation and size-exclusion chromatography. It is also possible that the potency of EVs is stronger in the presence of other ES factors or that the parasites contribute to potency of ES and EVs, for example by disrupting the mucus layer and enabling access to enteroid cells. It is also possible that ES products are dynamic and their content could differ when parasites sense the host epithelium. For example, *H. bakeri* ES from L4 larvae developing in colitis mice have altered secretomes than those from control mice suggesting that parasites can sense environmental changes and modify their ES accordingly ^44^.

Previous work has investigated the impact of adult HES on the basal epithelium using 3-D organoids^16,26^. We found that co-culture of EVdepHES, at doses comparable to others, with the apical surface of 2-D enteroids had subtle effects suggesting the basal epithelium is more responsive to foreign material, which is consistent with the basal localisation of key molecules involved in sensing PAMPs ^27,40^. Despite EVdepHES having subtle effects on the apical epithelium, the transcriptional changes were consistent with those induced by the live parasites in this study. Our results are also in agreement with the previous effects of HES treatment on 3-D organoids, including upregulation of genes related to epithelial repair (*Anxa8*, *F3, Fn1*) (although neither *Clu* nor *Ly6a* were significantly changed in our dataset); and the downregulation of *Clca1,* a gene that regulates mucus production by goblet cells in the lung and colon (a process key to anthelmintic defence) ^16,26,45^.

Using 2D-enteroids we observed lower levels of EV internalisation than those previously found by our group using the MODE-K cell line and bone marrow macrophages ^22,46^. This could be explained by features of the intestinal epithelium that are recapitulated by 2-D enteroids but missing in cell lines, which may impact *H. bakeri* EV internalisation. These features include production of mucus and secreted molecules, and the presence of cell subtypes that could have different specificities for EV uptake. Strikingly, EV-647+ cells displayed a strong upregulation of many ISGs (*Isg15*, *Cxcl10* and *Mx2)* and epithelial repair genes associated with revSCs. This could suggest that enteroid cells internalising EVs are revSCs, or that uptake of EVs induces differentiation of cells into this population. Further work is required to understand exactly how EVs act within these cells and whether they contribute to the induction of this cell population.

Here, we detected a strong ISG signature during the co-culture of worms with the basal epithelium and in cells that uptake *H. bakeri* EVs, but in the absence of host microbiota or hematopoietic immune cells, suggesting a direct ISG induction in response to *H. bakeri.* Indeed, *H. bakeri* infection has been associated with the expression of ISGs ^48,49^. Type I IFN responses are also important for effective immunity to parasitic worms, for example functional type I IFN receptor (*Ifnar*) is required for DCs to optimally prime CD4+ T cells during *Schistosoma mansoni* and *Nippostrongylus brasiliensis* infections ^50–52^. Further, *H. bakeri* infection of *Ifnar*^-/-^ mice suggests a role for type I IFNs in limiting worm fecundity and granuloma formation *in vivo,* however the contribution of the epithelium towards this is unknown ^12^. Interestingly, with the exception of *Ifn1b* which was detected at low levels (<1.8 CPM) only in sorted EV+ cells, no other type I or II (*Ifng)* IFN genes were detected by sequencing in this study. This agrees with our previous work where we found that infection of 2-D caecaloids with *T. muris* L1 larvae led to an expansion of a population of enterocytes expressing several ISGs (including *Isg15*), however, neither *Ifna* nor *Ifnb* was detected ^53^. This begs the question of how parasitic nematodes induce ISG expression in intestinal epithelium in an IFN-independent manner. Expression of several ISGs including *Isg15*, *Ifit1*, *Ifit2*, *Mx1* and *Viperin,* which are highly expressed in EV-647+ cells, can occur independently of IFNα/β through the detection of self (e.g., released during tissue damage) or foreign nucleic acids leading to IRF3 dependant expression ^54,55^. It is possible that *H. bakeri* RNA or DNA from live worms, either secreted as free extracellular RNA or transported by EVs in the case of EV+ cells, could initiate host expression of ISGs independent of type I IFNs.

In conclusion, 2-D enteroids provide an *in vitro* model that allows for co-culture of GI nematodes to closely recapitulate *in vivo* localisation to the epithelium. This model enables a better understanding of the direct contributions of parasite localisation, life-stage and isolated ES products on intestinal epithelial responses as well as the different responsiveness of the basal and apical sides of the epithelium. This model has also demonstrated the specificity of cellular populations for EV uptake and paves the way for more directed studies of EV functions and mechanisms in modulating specific cell types such as revSCs. In the future, 2-D organoid co-cultures can be modified to introduce other host factors such as immune cells, stromal cells and cytokines in a controlled manner to further replicate the parasite niche and allow for a holistic and mechanistic understanding of host helminth interactions during infection.

## Methods

### 3-D enteroid culture

3-D organoids were generated from C57BL/N6 mice (6-8 weeks) by dissecting the duodenum opening the intestine longitudinally and then cutting into ∼1-2 mm pieces. Tissue pieces were washed with Dulbecco’s PBS 1X without calcium and magnesium (PBS) (Sigma Aldrich, USA) using a pre-coated serological pipette until the supernatant was clear of intestinal contents and mucus (10-15 times) and incubated with Gentle Cell Dissociation Reagent (STEMCELL Tech, Canada) for 15 min at room temperature (RT) while gently rocking. Crypts were pelleted by centrifugation (1200 rpm, 3 min), washed with PBS, and seeded in 200 μl of Matrigel^®^ (Corning, USA) in 6-well cell culture plates and overlaid with organoid growth media containing organoid base growth medium (Advanced DMEM/F12, 1 x N-2 Supplement 1 x B-27 Supplement, 2 mM L-glutamine, 10 mM HEPES, (all from Gibco, Thermo Fisher Scientific, UK)) supplemented with 50% Wnt3a conditioned media (L-Wnt3a cells were kindly provided by from Hans the Clevers Laboratory, Utrecht, Netherlands), 10% R-spondin conditioned media (HA-R-spondin-1-Fc 293T cells (Amsbio, UK)), 1% Penicillin-Streptomycin (P/S, Gibco), 50 ng/ml Epidermal Growth Factor (EGF, Thermo Fisher Scientific), 1 mM N-acetylcysteine (NAC, Merck), 100 ng/ml recombinant Noggin (Peprotech, USA) and 10 μM Y-26732 dihydrochloride monohydrate (Rho Kinase inhibitor (ROCKi, Sigma-Aldrich, UK). Organoids were incubated at 37°C 5% CO_2_ and media was replenished every other day for 1 week after which ROCKi, Wnt3a CM and Penicillin/Streptomycin were removed.

### 2-D enteroid cultures

3-D organoids grown for 4 days in stem cell enrichment media comprising organoid growth media without Wnt3a CM and Penicillin/Streptomycin and supplemented with 10 μM CHIR99021 (Cayman Chemicals, UK)) were dissociated into a single cell suspension using TrypLE Express (Gibco). 200,000 cells in 200 μl of organoid base growth medium were seeded in a 12 mm Snapwell^TM^ cell culture insert with 0.4 μm pore (Corning) pre-coated with of Matrigel^®^ diluted 1:50 in ice-cold PBS. Basal compartments were supplied with 4 ml of stem cell enrichment media. 2-D organoids were incubated at 37°C, 5% CO_2._ and non-adherent cells were removed from the apical compartment and replaced with organoid base growth media after 1 day in culture and every second day onwards. Basolateral media was replaced every second day after initial seeding and organoids were grown for 4 days before use for characterisation, or co-culture with *H. bakeri* or ES products.

### Immunofluorescence microscopy

2-D enteroid apical and basal media was removed and the transwell membrane washed with PBS before fixation with 4% Paraformaldehyde (PFA) for 20 min at 4°C. PFA was removed, and the membrane washed 3 times by incubating with PBS for 5 min each at RT. If samples were treated with fluorescently labelled EVs they were covered from light throughout microscopy preparation. Samples were blocked with permeabilization/block buffer (1X PBS, 5% BSA, 2% Triton-X (Sigma)) for 1 hr at RT. Permaebilisation/block buffer was removed and samples were incubated with primary antibodies α-Lysozyme 1 (1:40, DAKO, A0099), α-DCAMKL1 (Abcam, UK, ab31704), α-Villin (1:100, Abcam, ab130751), α-Chromogranin-A (1:50, Abcam, ab15160), α-Ki67 (1:250, Abcam, ab16667) and the lectin dyes *Sambucus Nigra* FITC (SNA 1:50, Vector laboratories, UK, FL-1301) and *Ulex europaeus* FITC (UEA 1:1000, Sigma-aldrich, 19337) in diluent (PBS, 5% BSA, 0.25% Triton-X) overnight at 4°C. The following day antibodies were removed, and samples were washed 3 times with 200 μl diluent for 5 min at RT. Secondary antibodies α-Rabbit AF594 (1:1000, Biolegend, 410407), DAPI (Thermo Fisher Scientific) and Phalloidin-iFluor 594 (1:2000, Abcam) were added for 1hr at RT covered from light. Samples were washed 3 times with PBS. PBS was removed and the membrane was carefully excised using a scalpel, the Transwell^TM^ membrane was then placed apical side face up on a clean glass microscopy slide and 200 μl of Prolong Gold Antifade Mountant (Thermo Fisher Scientific). Fluorescence microscopy was performed using EVOS (Thermo Fisher Scientific). Super-resolution imaging was performed using the Zeiss LSM980 Airyscan 2, or the Zeiss LSM880 Airyscan microscope at the Centre Optical Instrument Laboratory (COIL) at the Wellcome Centre for Cell Biology, Edinburgh.

### Mice

C57BL/N6 & CBA 3 C57BL/6 F1 (CBF1) mice were housed at a temperature of 19–24 °C, with humidity between 40 and 65% under a 12-h light/dark cycle. To minimize suffering all efforts were made to provide considerate husbandry and housing and access to food (RM3 (P) irradiated diet) and water was provided *ad libitum*. Animal welfare was assessed routinely, and all mice were naïve before experimentation. Experiments were performed under the regulation of the UK Animals Scientific Procedures Act 1986 under the Project license P635073CF and experimental procedures approved by the University of Edinburgh Biological & Veterinary Services.

### *H. bakeri* lifecycle and HES generation

*H. bakeri* was maintained in CBF1 mice following established protocols ^56^. L4 larvae were isolated from the small intestine submucosa on day 7 by gentle extraction using forceps to burst the granuloma. Adults were collected at day 14 p.i. from the small intestine. Parasites were washed with Hank’s balanced salt solution (HBSS) with no calcium or magnesium (Sigma Aldrich) containing 1% P/S 6 times before soaking worms in 10% Gentamicin (Gibco) diluted in HBSS for 20 min and then washing 6 more times in HBSS containing P/S. Worms were cultured in serum-free RPMI 1640 supplemented with 1% D-Glucose (Merck),1% P/S, 1% L-glutamine (Gibco), and 100 μg/ml Gentamicin (Sigma Aldrich). After 24 h in culture, the media was replaced with fresh media to reduce the chance of contamination with host material. HES was collected from adult worms at days 4 and 8 in culture, eggs were pelleted by centrifugation at 400 x *g* (5 minutes) and HES supernatant passed through a 0.22 μm syringe filter (Merck) and stored at -80°C.

### Isolation and fluorescent labelling of extracellular vesicles

50-100 ml of HES was pooled and concentrated to <12.5 ml in 5 kDa MWCO Vivaspin^®^ (Sigma Aldrich) and EVs were isolated by ultracentrifugation (UC) at 100,000 x *g* for 2 h using a SW40 Ti rotor (Beckman Coulter) at 4°C. EV pellets were washed twice with 0.2 μm pre-filtered PBS and centrifuged as above but for 70 min. EV pellets were re-suspended in 100-200 μl PBS. EVs were then isolated by a secondary method of size exclusion chromatography (SEC) with in-house size exclusion columns of Sepharose CL-2B (Cytiva, USA). SEC fractions (0.5 mL) were collected and fractions 7-10 validated to contain EVs using the Zetaview twin laser (Particle Metrix, Germany). Fractions 7-10 were then combined and concentrated to ∼200 μL using a 5 kDa MWCO Vivaspin. For experiments with fluorescently labelled EVs, EVs were labelled after UC isolation by incubating with AlexaFluor647 Succinimidyl Ester (Thermo Fisher Scientific) for 1 h at RT. Excess dye was removed during the secondary isolation by SEC. To generate HES depleted of EVs (EVdepHES) supernatant was collected after the first UC of HES and concentrated to ∼1 ml. SEC was performed for further fractionation of EVdepHES and the fractions that do not contain EVs (fractions 8-30) were pooled and concentrated as above. Protein concentrations were quantified by Qubit (Thermo Fisher Scientific) and particle concentrations were calculated using Zetaview twin laser (Particle Metrix, Germany).

### Co-culture of *H. bakeri* parasites/ES products with 2-D enteroids

2-D enteroids were co-cultured with 20 μg/ml of EVs (which equates to an average of 2.6×10^10^ particles/ml) or 20 μg/ml EVdepHES in the apical compartment for 24 h. The concentrations were measured using Qubit 3.0 Fluorometer (Life Technologies). Live worm co-cultures were performed by incubating 10 adult worms or L4 larvae (isolated from infected mice at d7 p.i.) in the apical or basal compartment for 24 h.

### Dissociation of 2-D enteroids and Fluorescence Assisted Cell Sorting (FACS)

Parasites were removed carefully by pipette and then apical and basal media was aspirated from 2-D enteroids. The apical surface was washed three times with 1x PBS pre-warmed to 37°C. Single-cell suspensions from monolayers were prepared by three consecutive 10-minute incubations at 37°C in Trypsin 0.25% EDTA (Gibco) supplemented with 10 μM ROCKi. After each incubation, trypsin-containing cells were collected and washed three times in DMEM/F12 supplemented with 10% FBS and 10 μM ROCKi. Cells were pelleted at 200 x *g*, 4 min and re-suspended using wide-bore p1000 tips in 2 ml of TrypLE^TM^ Express Enzyme (Gibco) containing 10 μM ROCKi, 1 mM NAC (Merck) and 10 μg/ml DNase (STEMCELL Technologies) and incubated at 37°C in a water bath until a single cell suspension was observed (3-5 min). Single cells were pelleted by centrifugation and washed as above. Cells were resuspended in 1x PBS containing 10 μg/ml DNase and stained using LIVE/DEAD™ Fixable Aqua Dead Cell Stain Kit (Thermo Fisher Scientific, 1:1000) for 10 min protected from light at RT. Cells were washed twice in FACs buffer (1X PBS, 0.5% Bovine Serum Albumin (BSA), 2 mM EDTA (Thermo Fisher Scientific)). Sorting was performed using the FACS Aria (BD Biosciences) at 4 °C with a 70 μm nozzle at 60 psi.

### RNA extraction and sequencing

For bulk RNA sequencing, parasites, EVdepHES or EVs were removed from the culture and 2-D enteroids were washed three times in 1x PBS. Cells were lysed in transwells by adding 350 μl RTL lysis buffer to the membrane surface with pipetting to wash the membrane surface and collect lysate (Qiagen, Germany). For FACs experiments, cells were sorted directly into DNA LoBind^®^ Eppendorf tubes (Qiagen) pre-loaded with 350 μl RTL lysis buffer containing 3.5 μl of β-mercaptoethanol. In both cases cell lysates were homogenised immediately by vortexing for 1 min. RNA was extracted using RNeasy micro kit (Qiagen) as per kit instructions and RNA integrity (RIN) assessed by Bioanalyser or TapeStation (Aligent). The optional on-column DNA digest step was performed. For bulk 2-D enteroid experiments Ribodepletion, library preparation and sequencing were performed by the WT CRF using the NextSeq 200 P2 for 100 cycles. For RNA sequencing from sorted cells, cDNA amplification was performed using SMART-Seq^®^ v4 Ultra^®^ Low Input RNA Kit for Sequencing (TAKARA bio) and sequenced using NovaSeq SP flow cell with 50PE reads by Edinburgh Genomics.

### Analysis of RNA sequencing data

#### Quality control

Sequencing files were assessed for quality and filtered using FastQC, and low quality nucleotides and adapter sequences were removed using trimmomatic 0.39 with parameters ILLUMINACLIP:TruSeq3-PE-2.fa:2:30:10, LEADING:3, TRAILING:3, SLIDINGWINDOW:4:15 and MINLEN:36. Read alignment was performed using hisat2 version 2.2.1 using the non-deterministic parameter to the *mus musculus* genome from Ensembl release103 (genome GRCm39.primary_assembly.genome.fa). The resulting bam files were indexed using SAMtools (release 1.13) ^57,58^.

#### Read counts

Reads counts for each gene were generated within the package Rsubread 2.6.4 using the function featureCounts ^59,60^. The default parameters were used with the following modification to certain parameters, as follows, geneID was used for retrieving reads to generate counts. Multimapping reads were counted for all their reported mapping locations by setting countMutliMappingReads to true and fraction also set to true. Reads that overlapped multiple genome features were only counted for one genome feature and this was the feature with the largest number of bases aligned.

#### Differential expression analysis

Differential expression analysis (DEA) was performed using edgeR 3.34.1 on R version 4.1.1 ^60,61^. For pulling gene annotations (chromosome, gene symbol and description) biomaRt 2.48.3 was used. For co-culture with parasites or their ES products genes expressed with <1 CPM in at least 3 libraries were filtered out of the analysis. Tagwise dispersion estimates were performed using the function estimateDisp. Plots of DEA results were generated in R using the packages ComplexHeatmap and ggplot2.

#### CAMERA pathway analysis

Competitive gene set accounting for inter-gene correlation (CAMERA) was performed using all of the gene expression changes resulting from DEA analysis (no fold change or P-value cut-offs were used). CAMERA was performed in R using the limma package and the function camera ^38^. For CAMERA analysis using gene ontology, the gene ontology (GO) annotations were retrieved from org.Mm.eg.db 3.13.0. GO terms with fewer than 50 genes or more than 300 genes were removed from the analysis. For CAMERA analysis of cell type-specific genes, genes were retrieved from Haber *et al,* 2017 and/or Karo-Atar *et al*, 2022 publicly available single-cell RNA seq dataset of cells from *Mus musculus* intestinal epithelium or HES treated 3-D enteroids respectively ^16,37^.

## SUPPLEMENTARY FIGURES

**Figure S1:** Heatmap of all YAP/TAZ target genes after 24 h co-culture of *H. bakeri* L4 basal, adult basal or adult apical compared to mock. YAP/TAZ genes were previously defined based on up-regulation in YAP overexpressing organoids and downregulation in YAP KO organoids by Gregorieff *et al*, 2015^35^.

**Figure S2:** A&B) Confocal microscopy of 2-D enteroids treated for 24 h with EV-647 (20 μg/ml). EV-647 in yellow, Phallodin-iFluor 594 labels actin in violet and DAPI marks nuclei in cyan. A) Zoom of overlay region of EV uptake & individual channels. B) Orthogonal view of EV uptake in 2-D enteroid.

**Figure S3:** Heatmap of all YAP/TAZ target genes in sorted EV-467+ vs. Bystander, EV-647+ vs. Mock or Bystander vs. Mock. YAP/TAZ genes were previously defined based on upregulation in YAP overexpressing organoids and downregulation in YAP KO organoids by Gregorieff *et al*, 2015^35^.

**Movie S1:** Z-stack projection of 2-D enteroid topology. F-actin labelled with phalloidin (white), microvilli marker Villin AF647 (red), goblet cells labelled with UEA & SNA FITC (green), and nuclei are stained with DAPI (blue).

**Supplemental Table 1**

Table of differentially expressed genes (Log2FC 0.585, FDR < 0.05) from live parasite and ES co-culture conditions.

**Supplemental Table 2**

Table of differentially expressed genes (Log2FC 0.585, FDR < 0.05) from sorted 2-D enteroid cells.

## DATA AVAILABILITY

The bulk 2-D enteroid RNA-Seq data produced for this paper is available through NCBI’s Gene Expression Omnibus (GEO) under accession number GSE283214 (https://www.ncbi.nlm.nih.gov/geo/query/acc.cgi?acc=;GSE283214). RNA-seq of sorted enteroid cells produced for this paper is available through NCBI’s GEO under accession GSE283215 (https://www.ncbi.nlm.nih.gov/geo/query/acc.cgi?acc=;GSE283215).

## ACKNOWLEDGEMENTS

We thank Yvonne Harcus for assisting with generation of parasite excretory-secretory material and life cycle. Cell sorting was performed by the Flow Cytometry and Cell Sorting Facility in Ashworth, King’s Buildings at the University of Edinburgh. The facility is supported by funding from Wellcome and the University of Edinburgh. Library preparation and total RNA sequencing was performed by the Wellcome Trust Clinical Research Facility in Edinburgh (WT CRF). Low input library preparation of sorted cells and sequencing was performed by Edinburgh Genomics at the University of Edinburgh. RW was funded by the Darwin Trust of Edinburgh. This work was supported by ERC Consolidator Award 101002385 to AHB and the Rosetrees Trust grant M813 to RW and AHB. MADC was supported by a National Centre for the Replacement, Refinement and Reduction of Animals in Research (UK) David Sainsbury Fellowship (grant NC/P001521/1) and a Sir Henry Dale Fellowship jointly funded by the Wellcome Trust and the Royal Society (222546/Z/21/Z). For the purpose of Open Access, the author has applied a CC BY public copyright licence to any Author Accepted Manuscript version arising from this submission.

## AUTHORS’ CONTRIBUTIONS

Conceptualization AHB, RW & MADC; Data curation RW & JRBB; Formal analysis JRBB & RW; Funding Acquisition RW & AHB; Investigation RW & ER; Methodology RW & MADC; Project administration RW & AHB; Resources RW, AHB & MADC; Software JRBB; Supervision AHB & MADC; Visualisation RW; Writing-original draft RW, AHB & MADC; Writing-review & editing RW, AHB, & MAD

