## Supplementary figures and images for "2-D organoids demonstrate specificity in the interactions of parasitic nematodes and their secreted products at the basal or apical intestinal epithelium"

### Supplemental Figure 1

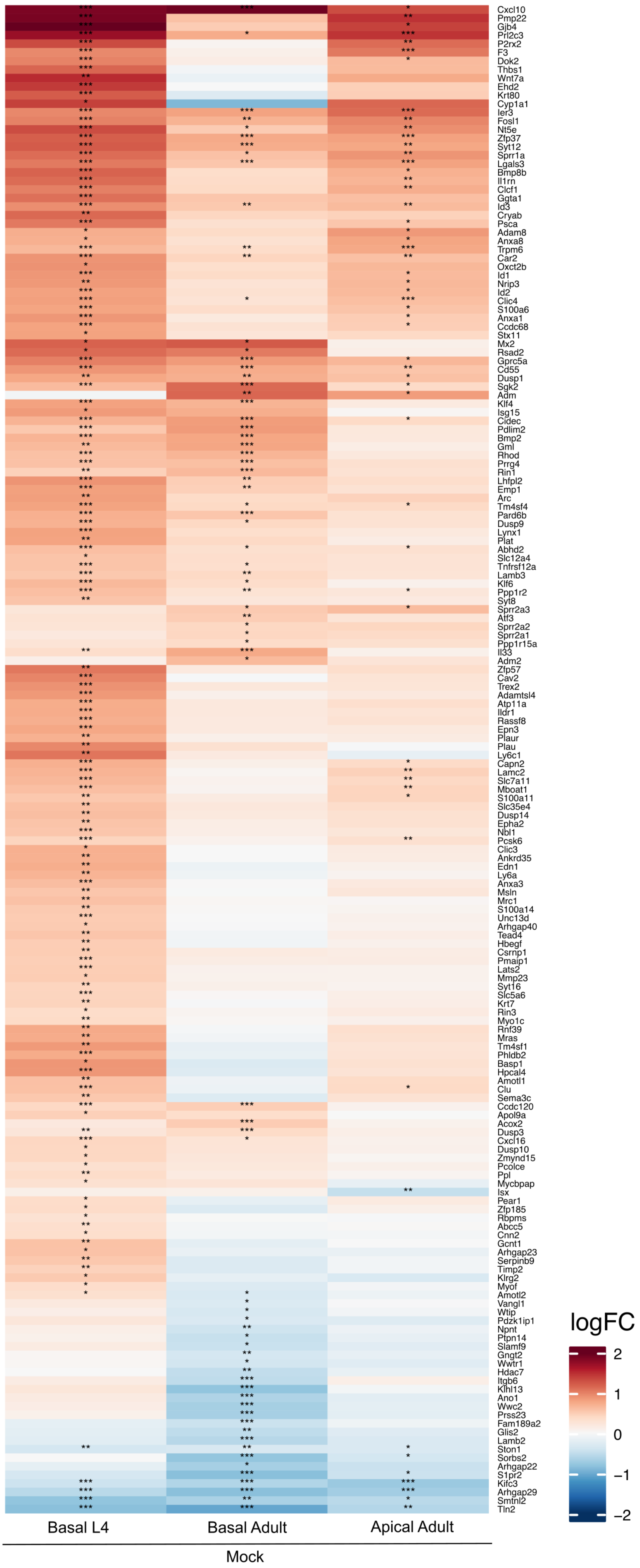

### Supplemental Figure 2

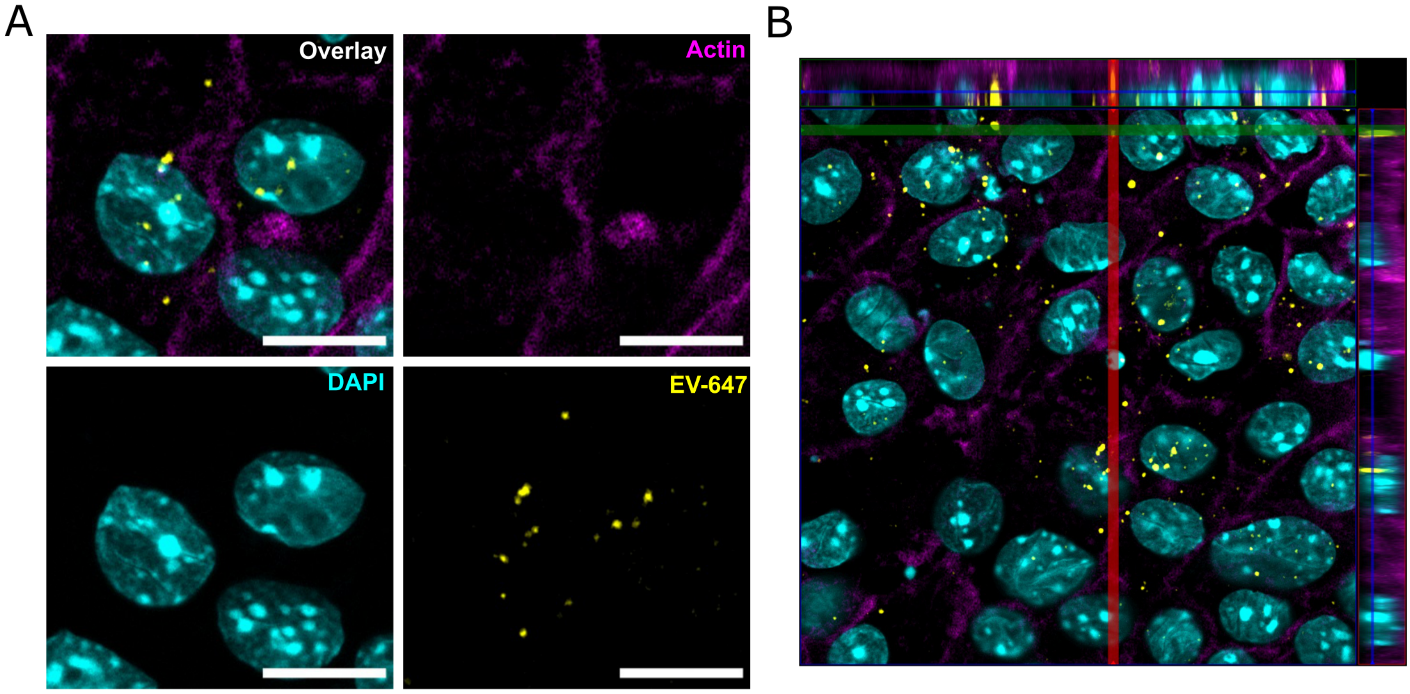

### Supplemental Figure 3

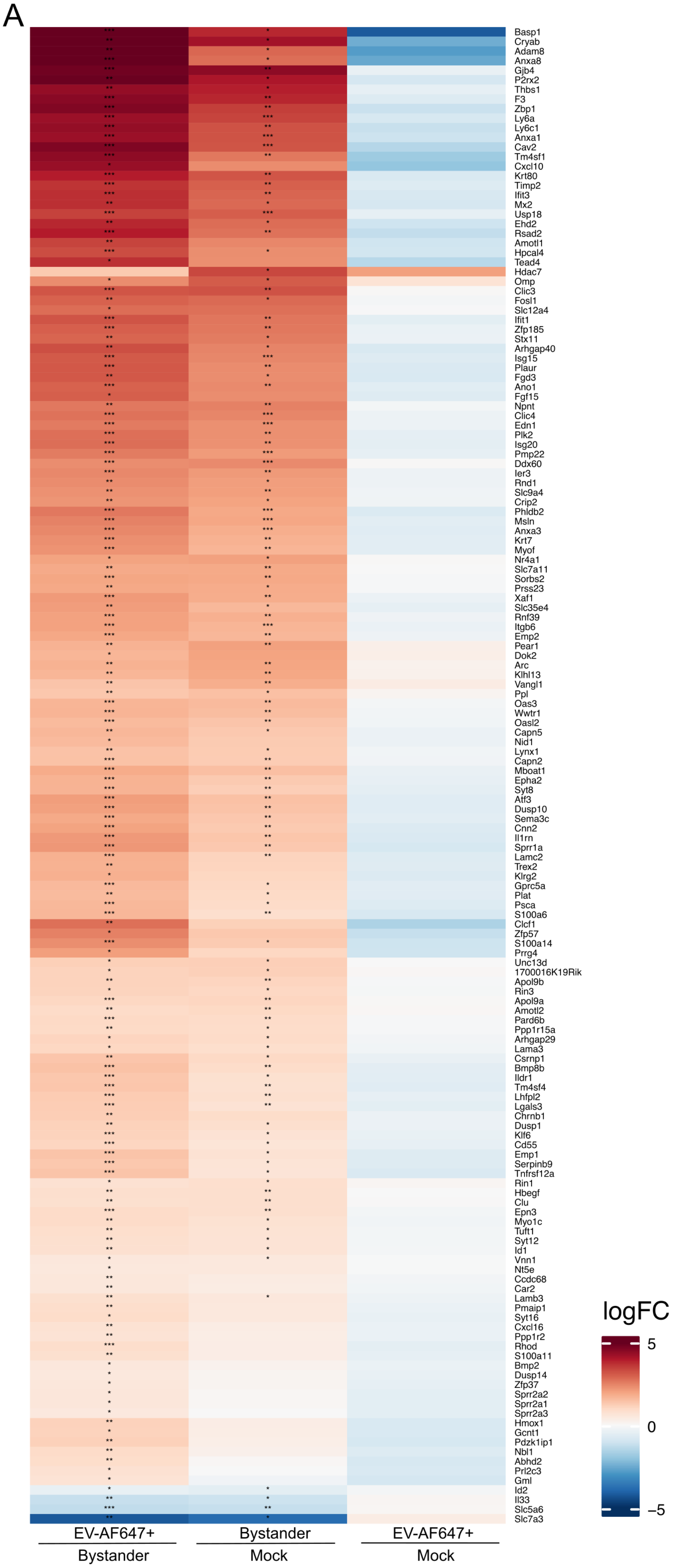
